# Water Stress Effect on the Growth of Lettuce (*Lactuca sativa*) and Moth Plant (*Araujia sericifera*)

**DOI:** 10.1101/2022.10.11.511498

**Authors:** Oluwatosin Adebanjo, Elikplim Aku Setordjie, Rong Wei Li

## Abstract

Water is the medium through which nutrients are transported from the soil into plants’ systems. Without soil water, the growth and yield of plants are negatively affected. This experiment compares the effect of water stress on the biomass production and chlorophyll content of lettuce (*Lactuca sativa*) and a weed species - moth plant (*Araujia sericifera).* The seedlings of the two plant species each were given three irrigation levels: 100% (T1), 50% (T2) and 0% (T3) field capacity of the growth medium - a mixture of peat moss (SuliFlor SF1^®^) and perlite (Perlindustria^®^) at ratio (3:1). The treatments for each plant were replicated five times, and the treatment lasted six weeks. Data were collected on moisture content and salinity of growing media, fresh and dry weight of shoot and root of plants, height, number of leaves, leaf area and chlorophyll content of leaf. The result showed that the water stress conditions have no significant effect on the chlorophyll, plant height and number of leaves of the two plant species. While the moth plant was not significantly affected by the stress conditions, lettuce recorded a significant reduction in leaf area, and in the dry weight of root and aerial part of the totally stressed plants, this shows that lettuce growth is significantly affected by water stress. Hence, moth plant tolerates water stress conditions more than the lettuce plant and this result may also be indicative of the survival of the moth plant if it infests a lettuce field.

## INTRODUCTION

In the wake of climate change, reports have been documented on the adverse impact of abiotic stress on the physiology and biomass production in many plants, resulting in up to 50% loss of agricultural output (Minhas *et al*., 2017). To curtail these stress conditions, plants have responded through different mechanisms and molecular networks (Osakabe *et al*., 2011; Ha et al., 2014). For example, in a stress condition associated with high solar radiation, some plants have induced a decrease in leaf water potential – shutting stomata opening or reducing leaf area to prevent water loss due to evapotranspiration (Osakabe and Osakabe, 2012). While the successful response of many plants to stress conditions is a function of the duration of the stress period (Osakabe et al., 2014), other plants have evolved genes that trigger the synthesis of compounds that help plants manage the stress condition (Vigeland *et al*., 2013).

Lettuce (*Lactuca sativa*) is an important temperate leaf vegetable, with Spain as the largest exporter and the second-largest producer after China (Aliste *et al*., 2020). Following the vitamin C content and polyphenols in lettuce, it has received so much attention in the human diet, such as in hamburgers, salads and a few other dishes. Its consumption has also been associated with weight loss (due to its low caloric content), lowered risk of cardiovascular diseases (via reducing low-density lipoprotein, cholesterol and blood pressure), and reduced risk of diabetes as it improves glucose metabolism (Antia *et al*., 2015). As with other crops, biotic and abiotic factors limit lettuce production. Although some abiotic factors such as water stress are known to induce the synthesis of phytochemicals with antioxidant properties which are of health benefit to humans (Myung-Min *et al*., 2010), they can also negatively affect the growth and productivity of lettuce, posing a threat to food security (Osakabe *et al*., 2014). This water stress condition can also be initiated by competition from weeds such as the Moth plant.

Moth plant (*Araujia sericifera* Brot.) is an invasive plant of southern Europe and the Mediterranean region (Coombs & Peter, 2010). It is native to South America and well-distributed, especially throughout southeast Latin America. Although cultivated as an ornamental plant in Italy and parts of the United States of America within the last century, it is recently considered a noxious weed, posing intense competition to crops (Di Noto and Castellano, 2010). As is characteristic of noxious plants, a wide range of mechanisms helps them survive stress conditions and to outcompete desired crops in the competition for soil water. Hence, this study compares the effect of varying water stress conditions on the growth of lettuce (*Lactuca sativa*) and moth plant (*Araujia sericifera*).

## MATERIALS AND METHODOLOGY

### 2.1. Experimental site

The trial was conducted in a greenhouse at the Polytechnic University of Valencia (39°29’00.5” N 0°20’30.0” W; 5 m A.S.L.), Spain, in autumn of 2021. The greenhouse structure is made of steel, with glasses on all sides with roof ventilation.

### 2.2. Plant material and growing condition

The seed of lettuce (*Lactuca sativa*) was purchased from Verdecora, Valencia, and planted for three weeks. Also, young wildings of moth plant (*Araujia sericifera*) were collected from the Puzol field in Valencia. Relatively uniform-sized lettuce seedlings and moth plant wildings were transplanted each into a pot of about 500 cm^3^ volume filled with growth media which is composed of peat (SuliFlor SF1 ^®^) and perlite (Perlindustria ^®^) at a ratio (3:1).

### 2.3. Experimental design

The two plant species were treated with three different levels of water stress; Control: 0 %, Intermediate stress: 50 % and Total stress: 100 % field capacity of the medium. The field capacity of the medium is estimated at 60 % moisture content. The experiment was laid out in a completely randomised design (CRD). The treatments were replicated five times, to make 30 experimental units.

For two weeks after transplanting, all plants received light irrigation before starting treatment for uniform stand establishment. The treatment lasted six weeks as plants were watered each time the moisture content of the control (T1) dropped to 50 per cent field capacity. This was done to maintain the irrigation levels.

### 2.4. Data collection

Before the transplanting, plant growth parameters such as plant height, number of leaves and leaf area were recorded. With the aid of W.E.T. sensor kit ^®^, each pot’s electrical conductivity (EC) and water content were measured. At six weeks after transplanting, all plants were harvested for the collection of data. Fresh root and shoot weight of plant were recorded using a weighing balance (g). Shoot and root were also oven-dried for two weeks at 60 °C fresh and weighed to get the dry root and shoot weight (g). Other records include leaf area (cm^2^) measured with the aid of Image J software, and number of leaf were also counted. The chlorophyll content of leaf was also measured using Konica Minolta ^®^ chlorophyll meter.

### 2.5. Data analysis

The data were analysed using R version 4.1.2. One-way ANOVA was conducted to determine the effect of the different treatments on the growth parameters of the two plants separately. As a posthoc test, Tukey’s HSD test was employed to evaluate the differences between the means of the different treatments per plant. Correlation analysis was conducted to determine the relationship between growing media parameters and the growth parameters of both plants used in the study.

## RESULTS

### 3.1. Progression of Moisture Content of Growing Media

Moisture content decreased in all treatments across both plants over the course of the experiment.

### 3.2. Progression of Salinity of Growing Media

Salinity increased significantly in most treatments across both plants over the course of the experiment except in the Control treatment (0% stress) of *Lactuca sativa* as indicated in Figure 3.

**Figure 1:**
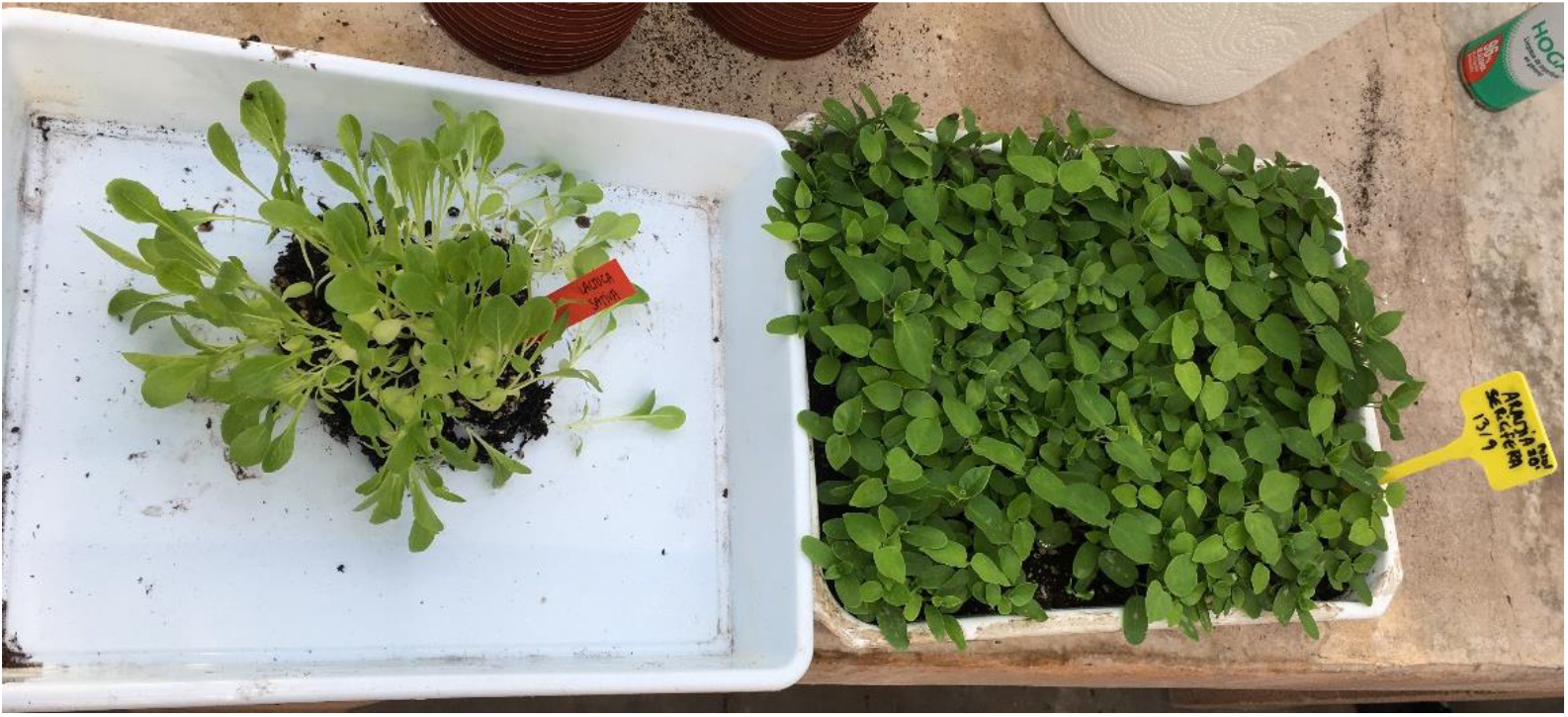
(Left to right) Seedlings of *Lactuca sativa* and *Araujia sericifera* just before transplanting

**Figure 2:**
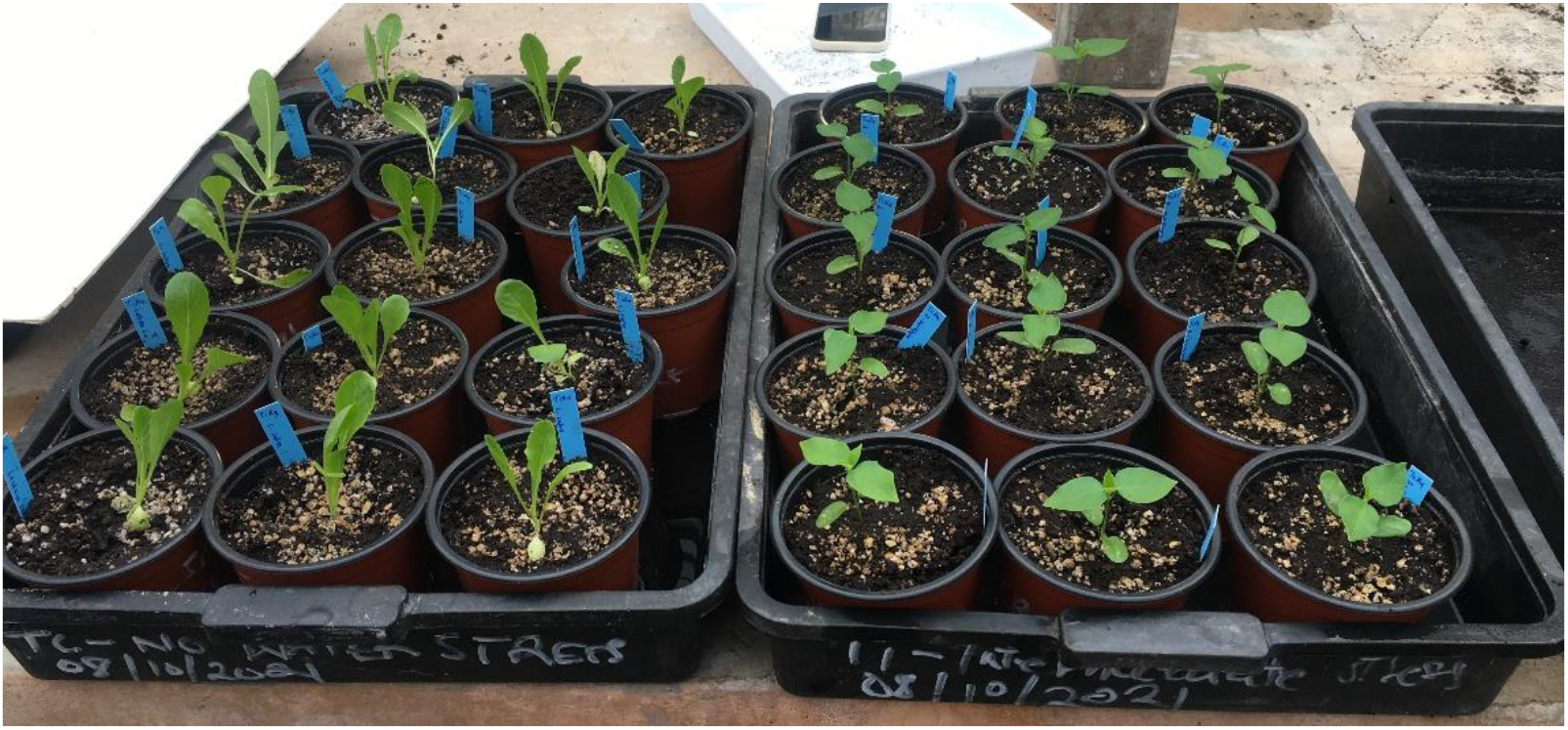
(Left to right) Seedlings of *Lactuca sativa* and *Araujia sericifera* just after transplanting

**Figure 3:**
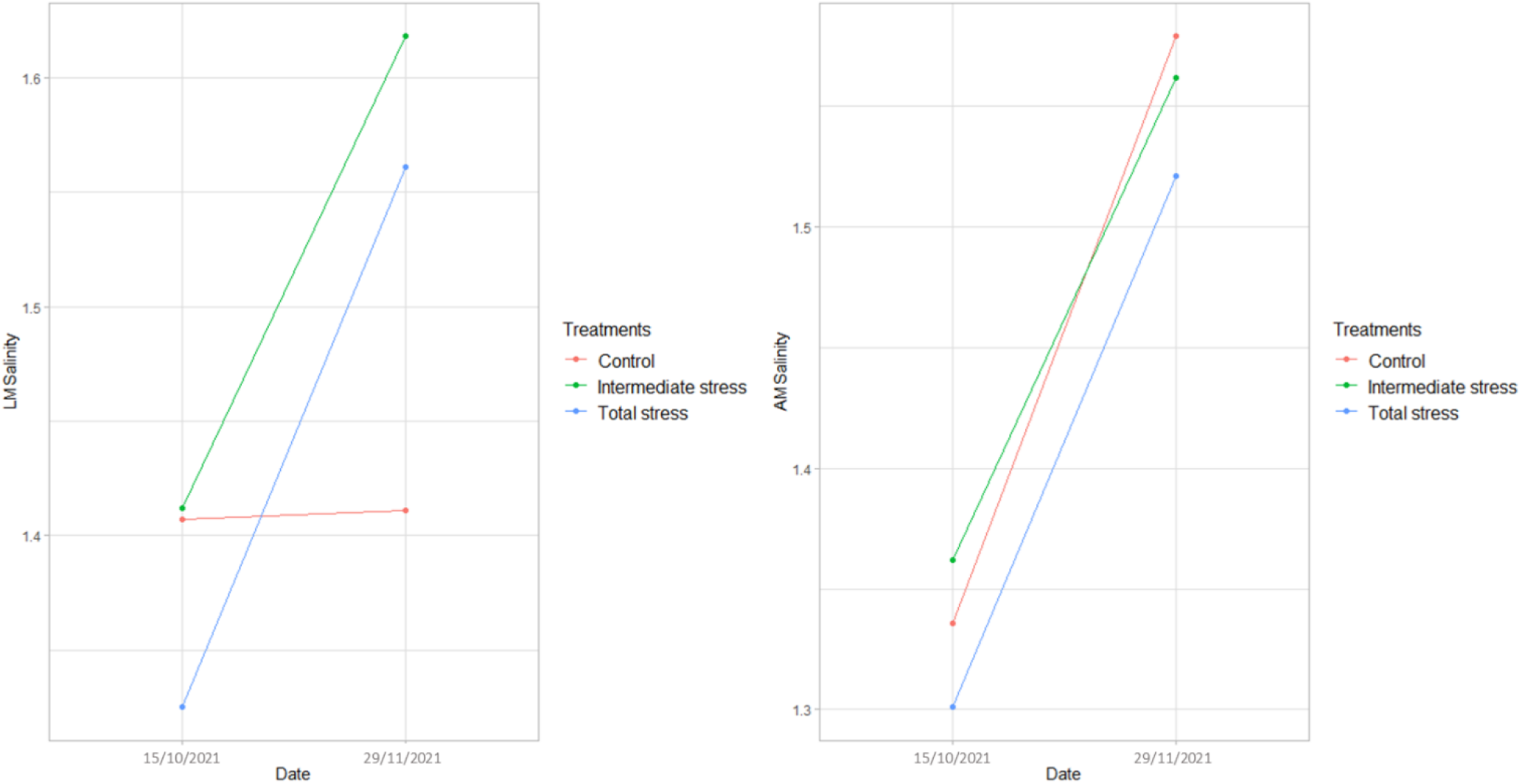
Trend of Salinity of Growing Media of *Lactuca sativa* (left) and *Araujia sericifera* (right) over time (control, intermediate stress and total stress denote water stress levels of 0, 50 and 100%, respectively).

### 3.3. Effect of Different Stress Levels on Moisture Content of Growing Media

The different treatments had no significant effect on the final moisture content (FMC) of growing media of *Lactuca sativa.* However, in *Araujia sericifera,* significant differences were observed. The highest moisture content was observed in Intermediate stress with a value of 28.81% followed by Total stress and Control with values of 22.78% and 16.60%, respectively (Table 1).

**Table 1.** Effect of different treatments on the final moisture content of both plants.

| Treatments | *Lactuca sativa* FMC (%) | *Araujia sericifera* FMC (%) |
| --- | --- | --- |
| **Control** | 11.11 a | 16.60 b |
| **Intermediate Stress** | 16.69 a | 28.81 a |
| **Total Stress** | 15.94 a | 22.78 ab |

\* Means followed by the same letters do not differ significantly at $p = 0.05$ .

### 3.4. Effect of Different Levels of Water Stress on the Growth Parameters

**Leaf Area:** Leaf area in *Lactuca sativa* in the different treatments were significantly different from each other, the highest leaf area was observed in *Lactuca sativa* plants subjected to Intermediate stress (142.3 cm^2^), followed by Control (114.4 cm^2^) and Total stress (94.4 cm^2^). Leaf Area was however not significantly different in the means of the three treatments in *Araujia sericifera* (Figure 4).

**Figure 4:**
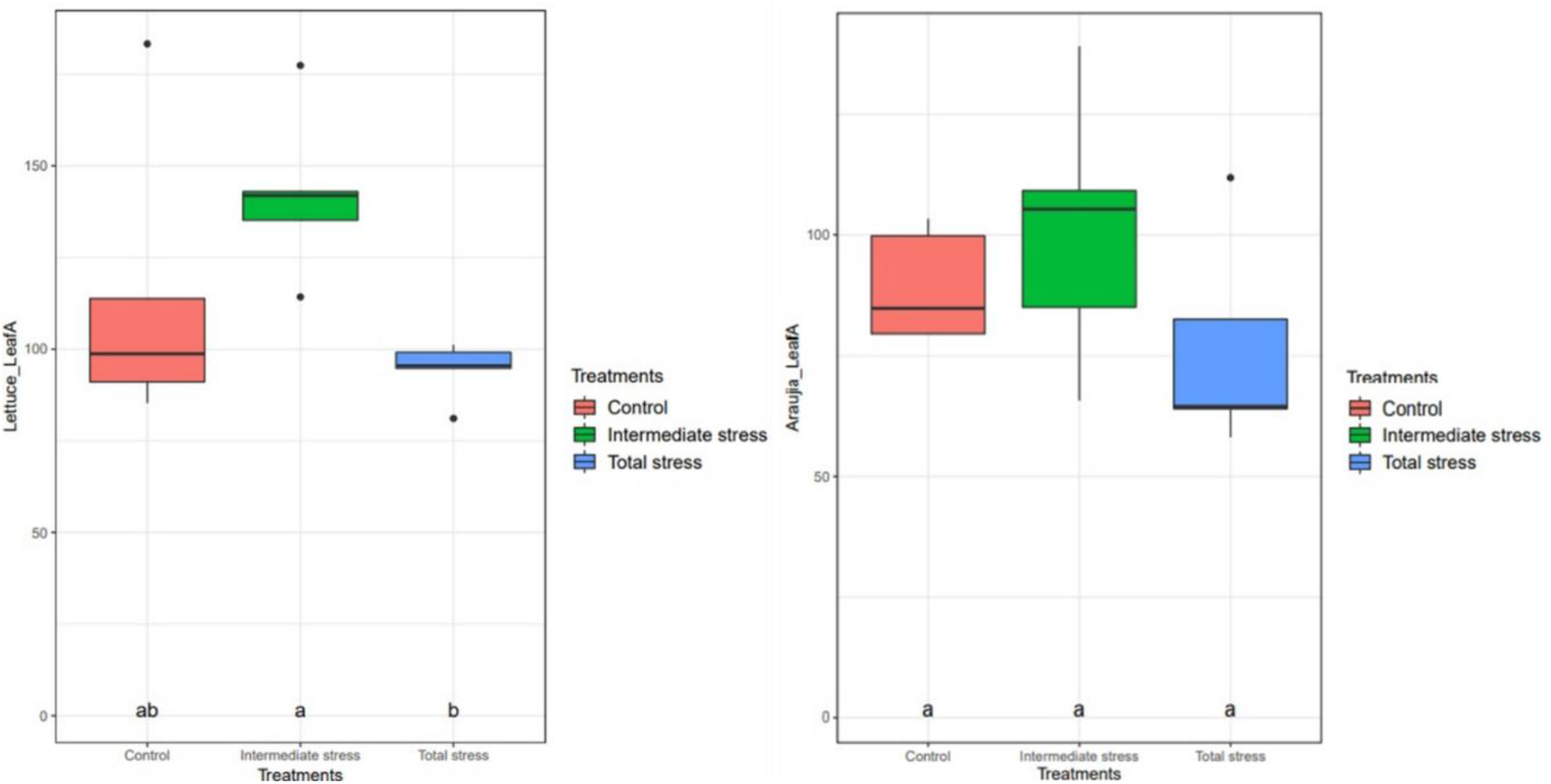
Effect of different levels of water stress on Leaf Area of *Lactuca sativa* (left) and *Araujia sericifera* (right) over time (control, intermediate stress and total stress denote water stress levels of 0, 50 and 100 %, respectively). Means followed by the same letters do not differ significantly at *p* = 0.05.

**Root Length:** Root length was significantly lower in the Control treatment (6.06 cm) of *Lactuca sativa* as compared to Intermediate Stress (13.48 cm) and Total Stress (13. 20 cm). There were no significant differences between treatments of *Araujia sericifera* (Figure 5).

**Figure 5:**
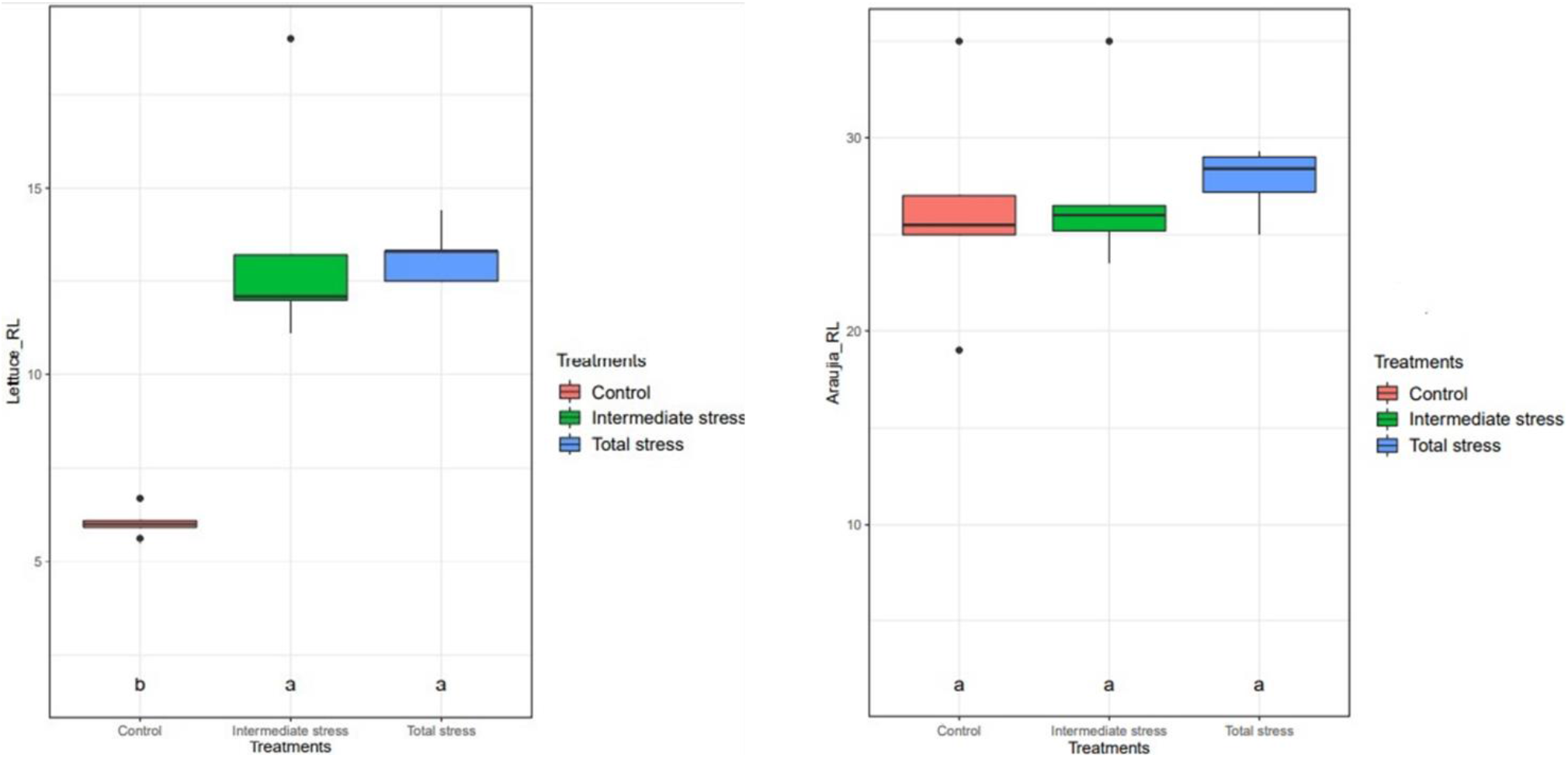
Effect of Different Stress Water Levels on the Root Length of *Lactuca sativa* (left) and *Araujia sericifera* (right) over time (control, intermediate stress and total stress denote water stress levels of 0, 50 and 100%, respectively). * Means followed by the same letters do not differ significantly at *p* = 0.05.

There were no significant differences between treatments in other growth parameters (Leaf number, Plant Height, Chlorophyll content, etc.) in both *Lactuca sativa* and *Araujia sericifera.*

### 3.5. Effect of Different Levels of Water Stress on Biomass

There were no significant differences in the Fresh Weight of all treatments in both plants. The Dry weight of Aerial parts was generally lower in *Araujia sericifera* than in *Lactuca sativa*. Dry Aerial Weight (DAW) and Dry Root Weight (DRW) in *Araujia sericifera* were generally lower than in *Lactuca sativa* however there was no significant difference between treatments in *Araujia sericifera.* DAW was significantly lower in the Total stress treatment of *Lactuca sativa* with a mean value of 0.816 g than Control and Intermediate stress treatments with mean values of 2.192 g and 3.036 g, respectively (Figure 6).

**Figure 6:**
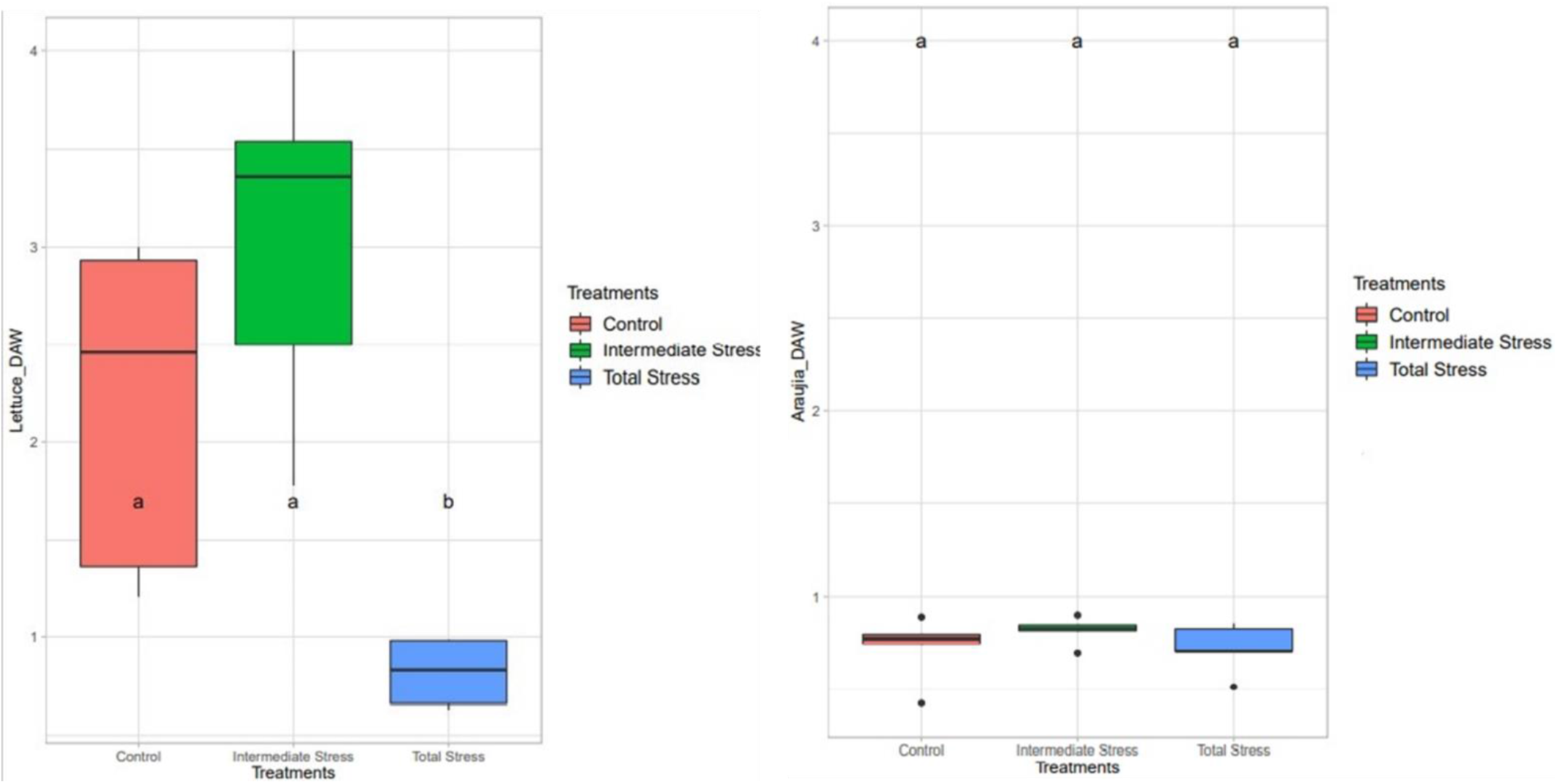
Effect of different levels of water stress on Dry Aerial Weight of *Lactuca sativa* (left) and *Araujia sericifera* (right) over time (control, intermediate stress and total stress denote water stress levels of 0, 50 and 100 %, respectively). * Means followed by the same letters do not differ significantly at *p* = 0.05

Significant differences were also observed in Dry root weight between all treatments of *Lactuca sativa.* Total stress (0.266 g) had the lowest Dry root weight followed by Control (0.694 g) and Intermediate stress (1.242 g) as observed in Figure 7.

**Figure 7:**
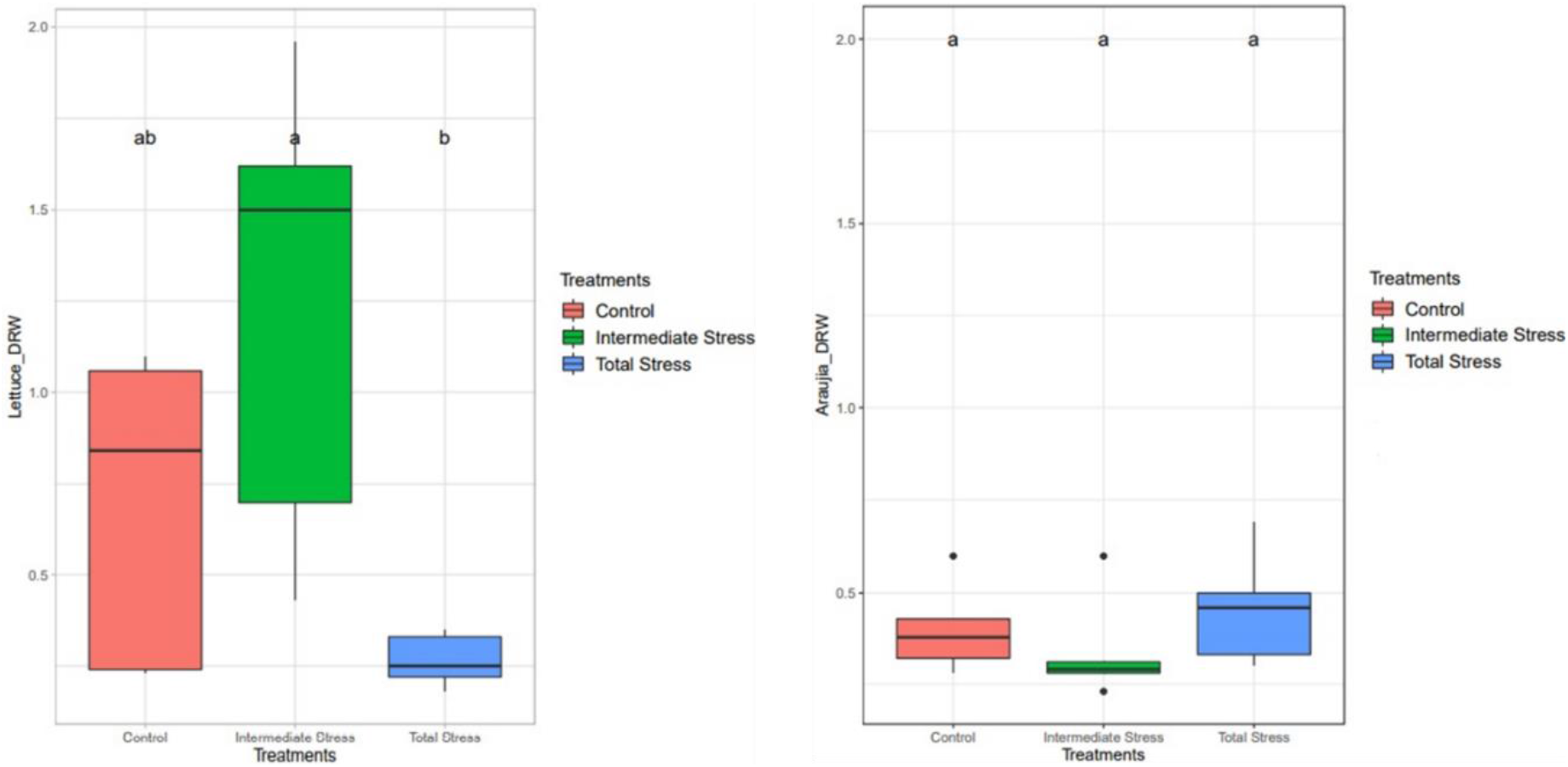
Effect of different levels of water stress on Dry Root Weight of *Lactuca sativa* (left) and *Araujia sericifera* (right) over time (control, intermediate stress and total stress denote water stress levels of 0, 50 and 100 %, respectively). * Means followed by the same letters do not differ significantly at *p* = 0.05

**Figure 8:**
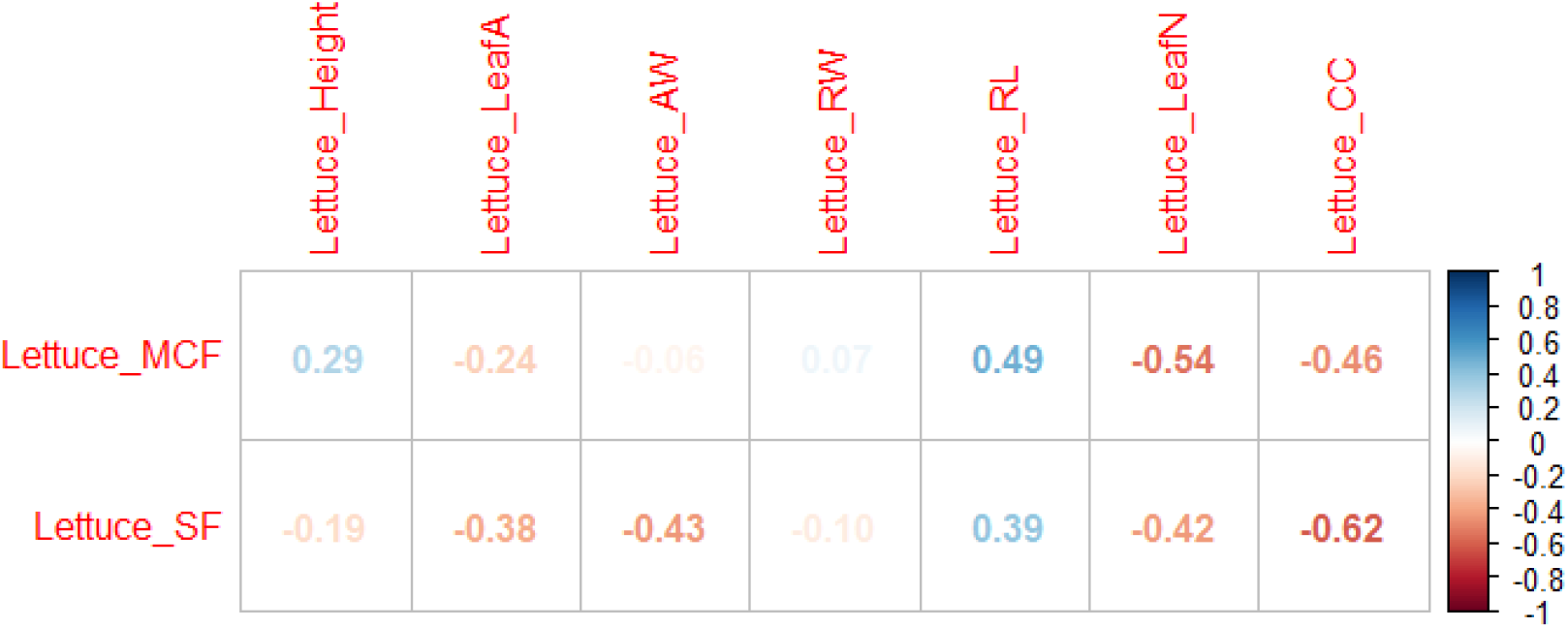
Pearson’s Correlation between Final Moisture Content (Lettuce_MCF), Final Salinity (Lettuce_SF) and Growth Parameters of *Lactuca sativa.* Showing the relationship between Final Moisture Content and Final Salinity to different growth parameters.

**Figure 9:**
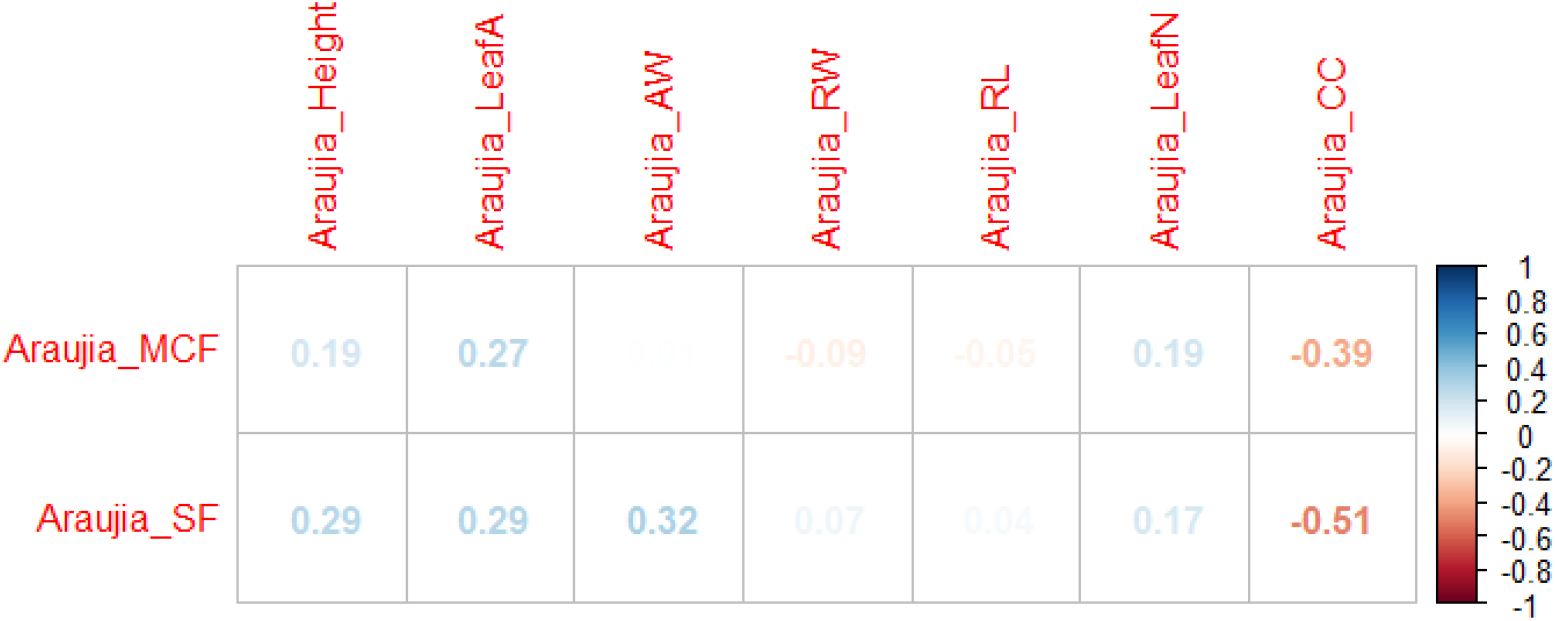
Pearson’s Correlation Values between Final Moisture Content (Araujia_MCF), Final Salinity (Araujia_SF) and Growth Parameters of *Araujia sericifera.* Showing the relationship between Final Moisture Content and Final Salinity to different growth parameters

## DISCUSSION

Plants absorb more water as they grow, hence the decrease of moisture content across all treatments in both plants over time. In relation, salinity increased as less water was available to flush out possible soluble salts or dissolved ions contained in the tap water used for irrigation (Maraver *et al*., 2015). This trend is however different in the Control of *Lactuca sativa,* where salinity increase was very low over the period of the experiment, and this could be attributed to optimal irrigation in this treatment.

Plant leaf area decreases with water stress, a similar trend observed in the study by Kurunc (2021), which indicated that increased water stress could lead to leaf curling, wilting and senescence. Our results showed the lowest leaf area in the Total stress treatment of *Lactuca sativa* in support of this trend. However, the Intermediate stress treatments had the highest leaf area.

Root Length in *Lactuca sativa* seedlings was significantly higher in the Intermediate stress and Total stress treatments, this phenomenon can be attributed to the rapid growth of plant roots towards the water table in drought conditions. This supports the finding of Gupta *et al.* (2020) on the rapid gravitropism of plant root in an intermediate drought condition.

Dry Aerial Weight and Dry Root Weight was significantly lower in Total Stress treatments of *Lactuca sativa,* this confirms research conducted by Kizil *et al.* (2012) which observed that *Lactuca sativa* yield is affected by low irrigation levels. Water stress is also known to have a destructive effect on root biomass.

According to Bellache *et al*. (2022), *Araujia sericifera* has a high tolerance to water stress. This explains the general lack of significant differences between the means of growth parameters in all three treatments of this plant.

*Lactuca sativa* seedlings suffered from pest and disease attacks during the experiment which also could have affected growth parameters.

## CONCLUSION

The results presented here show that water stress affects plant growth negatively. Although not so visible in growth parameters, data on dry plants clearly indicated the effect of water stress on *Lactuca sativa,* with Total Stress having the lowest dry weight compared to the other treatments. *Lactuca sativa* is more susceptible to water stress than *Araujia sericifera*. This result may also be indicative of the survival of the moth plant if it infests a lettuce field as it is may is not affected by the competition for water that may exist between both plants.

## Supporting information

Supplementary raw data

## Abbreviations

LeafA: Leaf Area;
HF: Final Height;
AW: Aerial Weight;
RW: Root Weight;
RL: Root Length;
LeafN: Number of leaves;
CC: Chlorophyll content;
DAW: Dry Aerial Weight;
DRW: Dry Root Weight

## AUTHORS’ CONTRIBUTION

Rong Wei Li used Image J software to obtain data for area of leaf. Elikplim Aku Setordjie organized and analysed the data, wrote the result and discussions. Oluwatosin Adebanjo wrote the abstract, introduction, materials and methods.

## CONFLICT OF INTEREST

The authors declare no conflict of interest is acceptable

## Notes

### Competing Interest Statement

The authors have declared no competing interest.

